# Enhancement of gene expression noise due to nonspecific transcription factor binding

**DOI:** 10.1101/2019.12.24.887984

**Authors:** Supravat Dey, Mohammad Soltani, Abhyudai Singh

## Abstract

The genome contains several high-affinity non-functional binding sites for transcription factors (TFs) creating a hidden and unexplored layer of gene regulation. We investigate the role of such “decoy sites” in controlling noise (random fluctuations) in the level of a TF that is synthesized in stochastic bursts. Prior studies have assumed that decoy-bound TFs are protected from degradation, and in this case decoys function to buffer noise. Relaxing this assumption to consider arbitrary degradation rates for both bound/unbound TF states, we find rich noise behaviors. For low-affinity decoys, noise in the level of unbound TF always monotonically decreases to the Poisson limit with increasing decoy numbers. In contrast, for high affinity decoys, noise levels first increase with increasing decoy numbers, before decreasing back to the Poisson limit. Interestingly, while protection of bound TFs from degradation slows the time-scale of fluctuations in the unbound TF levels, decay of bounds TFs leads to faster fluctuations and smaller noise propagation to downstream target proteins. In summary, our analysis reveals stochastic dynamics emerging from nonspecific binding of TFs, and highlight the dual role of decoys as attenuators or amplifiers of gene expression noise depending on their binding affinity and stability of the bound TF.

## Introduction

The level of a gene product can show remarkable cell-to-cell differences within an isogenic cell population exposed to the same external environment.^1–9^ This intercellular variation has been referred to as gene expression noise and is attributed to biochemical processes operating with low-copy number components, such as, the number of promoters and mRNAs for a given gene. Noise in expression critically impacts the functioning of diverse cellular processes in both beneficial and detrimental ways. For example, on one hand noise has been shown to be detrimental for developing embryos,^7,10,11^ organismal fitness^12^ and has been connected to disease phenotypes.^13,14^ On the other hand, noisy expression drives phenotypic heterogeneity between otherwise genetically-identical cells, allowing them to hedge their bets against environmental uncertainties.^5,8,15–24^ While there has been considerable work uncovering the role of transcription/translation processes together with molecular feedbacks in driving expression noise,^25–28^ how nonspecific binding of a protein shapes this noise remains poorly understood.

A fundamental step in gene regulation is the binding of a transcription factor (TF) to its target promoter.^29–31^ Besides this specific binding, TFs also bind to other non-functional high-affinity sites spread across the genome. These spurious binding sites, known as transcription factor decoys, can be present in different abundances with various binding affinities.^32–35^ The regularity role of decoys through molecular sequestration of TFs has been experimentally demonstrated via synthetic circuits in *Escherichia coli*^36^ and *Saccharomyces cerevisiae*,^37^ and not surprisingly, the TF-inhibiting activity of decoys has tremendous therapeutic potential.^38–41^ An immediate consequence of TF sequestration by decoys is the modulation of TF’s stability. Depending on the context, decoy binding can either enhance or reduce the stability of the TFs. For certain TFs such as MyoD, the DNA binding provide increased stability against degradation.^42,43^ On the other hand, for VP16 in *Saccharomyces cerevisiae*, the bound TFs become more prone to degradation by the ubiquitin-mediated proteolysis.^44^ A key focus of this work is to investigate how the relative stability of bound vs. unbound TF affects both the extent and time-scale of fluctuations in the level of a given TF.

Prior theoretical work of on this topic has highlighted the role of decoys as noise buffers, in the sense that, the presence of decoys attenuates random fluctuations in number of freely (unbound) available copies of the TF.^45–55^ However, these results are based on the assumption that sequestration of TF at a decoy site protects the TF from degradation. Relaxing this assumption to consider an arbitrary decay rate of the bound TF, we uncover a novel role of decoys as both noise amplifiers and buffers. We systematically characterize the parameter regimes leading to these contrasting roles in terms of the bound vs. free-TF stability, number and affinity of decoy sites. Finally, we study noise transmission from the TF to a downstream target protein reporting counterintuitive effects in some cases, for example, decoys amplify noise in the level of a TF but reduces noise in the level of the TF’s downstream target protein.

## Results

### Model formulation

To study the role of decoy binding sites, we consider a TF that is synthesized in stochastic bursts (Fig. 1). Such bursty expression of gene products has been experimentally observed in diverse cell types,^56–63^ and corresponds to distinct mechanisms at the transcriptional and translation level. For example, a promoter randomly switches from a transcriptionally inactive to an active state, and quickly turns off after producing a bursts of mRNAs.^59,64–66^ Moreover, assuming short-lived mRNAs, each synthesized mRNA is rapidly degraded after translating a burst of proteins.^67–69^ We phenomenologically model the combined effect of transcriptional and translational bursts by considering a Poisson arrival of burst events with rate *k_x_*, and each event creates *B_x_* molecules, where *B_x_* is an independent and identically distributed random variable following an arbitrary positively-valued distribution.^70–75^ More specifically, *B_x_* = *i* with probability *α_x_*(*i*) where *i* ∈ {1,2,…}. We consider a general form for the burst size distribution *α_x_*(*i*) throughout the paper, except for plotting and simulation purposes where *α_x_*(*i*) follows a geometrical distribution.^69,76–78^

**Figure 1.**
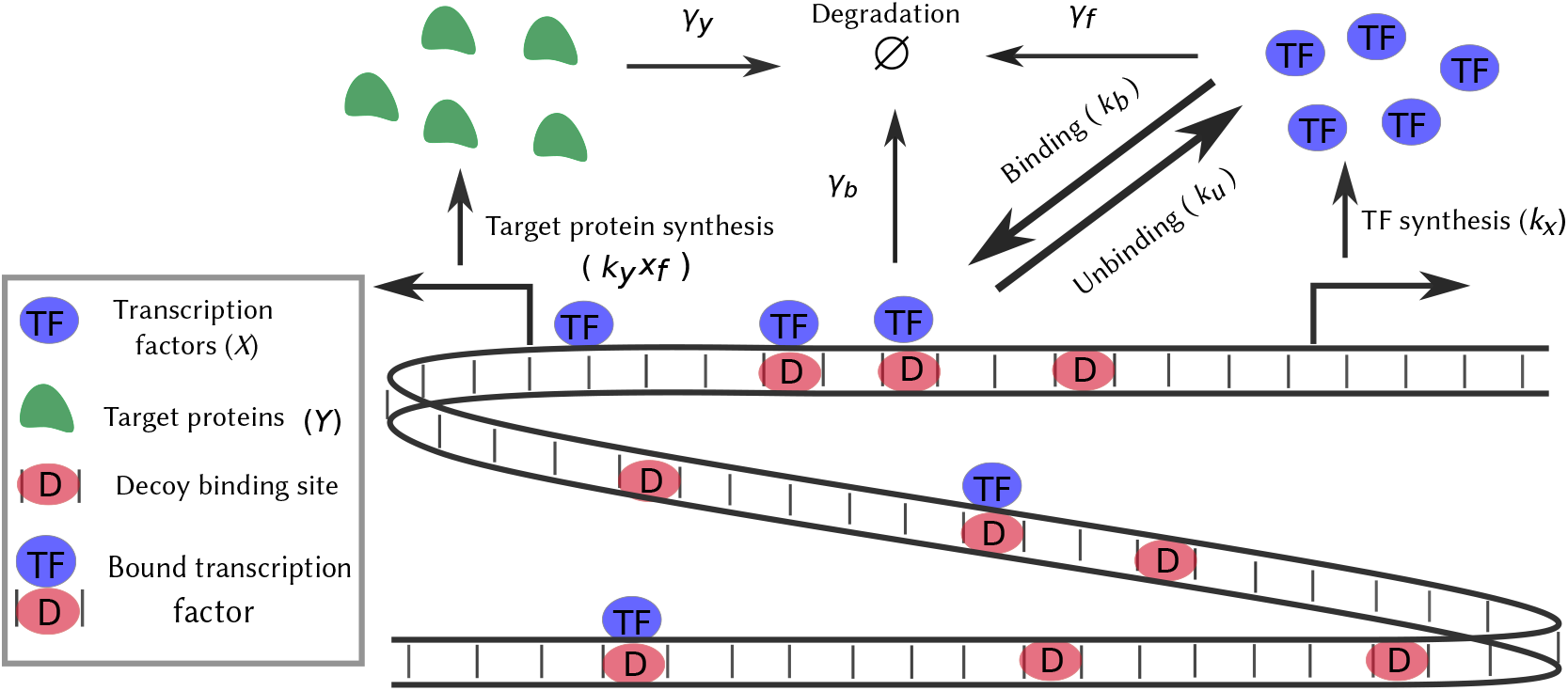
Model schematic for investigating the impact of nonspecific transcription factor binding on expression noise. A genome with several decoy binding sites where transcription factors (TFs) bind reversibly, is depicted. The synthesis of TFs occurs in stochastic bursts. Both the free and bound TFs are subject to degradation. The free TFs activate a target gene and regulate its bursty protein synthesis.

Consider *N* decoy binding sites in the genome, with the TF binding/unbinding to each decoy site with rates are *k_b_* and *k_u_*, respectively. As motivated earlier, we allow for both the free (unbound) and bound TF to decay with rates *γ_f_* and *γ_b_*, respectively. Let *x_f_*(*t*) denote the number of free TF molecules, and *x_b_*(*t*) the number of bound TFs at time *t* inside a single cell. Then, the stochastic evolution of *x_f_*(*t*) and *x_b_* (*t*) is governed by the following probabilities

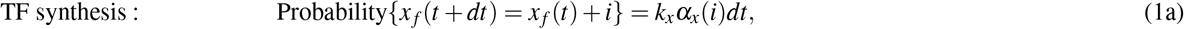

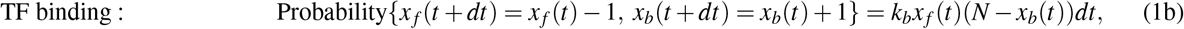

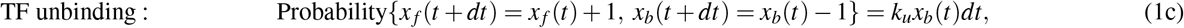

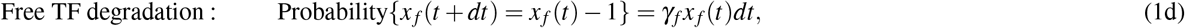

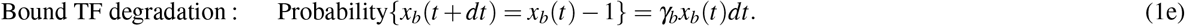

Each equation here defines a stochastic events that occurs with a certain probability in the small time interval (*t*, *t* + *dt*], and whenever the event occurs the population counts change by discrete integer amounts. Based on the underlying stochastic chemical kinetics, these occurence probabilities are either independent (as in 1a), or linearly/nonlinearly dependent on the molecular counts. For readers convenience, we provide a list of model parameters along with their description in Table 1. Expressing *γ_b_* in terms of *γ_f_* as *γ_b_* = *βγ_f_*, *β* = 0 corresponds to no decay of bound TF, and *β* = 1 corresponds to equal degradation rates for both the free and bound TF. The key focus of this work is to understand the stochastic dynamics of *x_f_*(*t*) in different regimes of *β*.

**Table 1.** Summary of notation used

| Parameter | Description | Parameter | Description |
| --- | --- | --- | --- |
| $x_f$ | Free TF count | $x_b$ | Bound TF count |
| $N$ | Total decoy binding sites | $k_x$ | TF burst frequency |
| $k_b$ | TF binding rate | $k_u$ | TF unbinding rate |
| $\gamma_b$ | Bound TF degradation rate | $\gamma_f$ | Free TF degradation rate |
| $\beta$ | $\gamma_b/\gamma_f$ | $k_d$ | Dissociation constant ($k_u/k_b$) |
| $y$ | Target protein count | $k_y x_f$ | Target protein burst frequency |
| $\gamma_y$ | Target protein degradation rate | $\alpha_x(\alpha_y)$ | Burst size distribution of $X(Y)$ |
| $\langle B_x \rangle (\langle B_y \rangle)$ | Average burst size for $X(Y)$ | $\langle B_x^2 \rangle (\langle B_y^2 \rangle)$ | Second moment of burst size distribution for $X(Y)$ |

Based on the above discrete-state continuous-time Markov model, one can write a corresponding Chemical Master Equation (CME) that provides the time evolution of the joint probability density function *p*(*x_f_*, *x_b_*, *t*), for observing *x_f_* free TF, and *x_b_* bound TF molecules at time *t*^79,80^

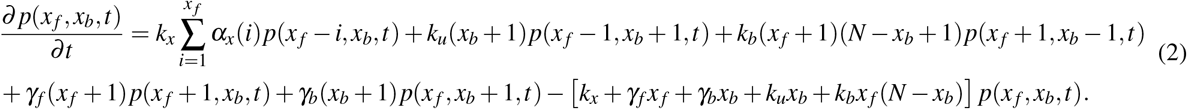

As the CME is analytically intractable, it is typically solved numerically through either the Finite State Projection algorithm^81–83^ or various Monte Carlo simulation techniques.^84–88^ Taking an alternative approach, we focus on the statistical moments of the molecular counts and use the well-known Linear Noise Approximation^89–94^ to obtain closed-form formulas for the mean and noise levels.

### Noise in free TF counts in the absence of decoy sites

When there are no decoys (*N* = 0), the moments of the free TF count can be solved exactly from the CME. In particular, the steady-state mean level 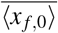, and the Fano factor *F*_*x_f_*,0_ (variance/mean) of *x_f_*(*t*) are given by,^26,95^

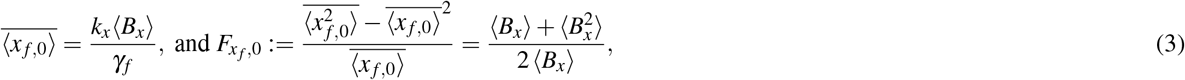

respectively, where 〈*B_x_*〉 and 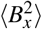 are the first and second-order moments of the burst size *B_x_*. Throughout the paper we use angular brackets 〈〉 to denote the expected value operation, and 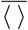 to represent the steady-state expected value. Note that the Fano factor is completely determined by the burst size distribution, and is independent of the burst arrival rate and the protein decay rate. As expected, we recover the Poisson limit (*F*_*x_f_*,0_ = 1) for non-bursty production *B_x_* = 1 with probability one, and noise is super-Poissonian (*F*_*x_f_*,0_ > 1) for bursty production. In the special case where *B_x_* follows a geometric distribution

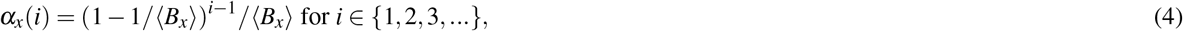

the steady-state Fano factor *F*_*x_f_*,0_ = 〈*B_x_*〉 is equal to the mean burst size.

### Bound TF’s degradation titrates the regulating activity of TF

In the presence of decoy (*N* > 0), exact solutions to the mean and noise levels are unavailable, and one has to resort to approximation techniques. Recall from (1), that the probability of the TF binding event is nonlinearly related to the molecular counts via the product term *x_f_*(*t*)*x_b_*(*t*). This nonlinearity results in unclosed moment dynamics – the time evolution of lower-order moments depends on higher-order moments^96,97^. For example, the dynamics of the means 〈*x_f_*〉, 〈*x_b_*〉 depends on the second order moment 〈*x_f_x_b_*〉 (see SI, section 1). Typically approximate closure schemes are employed to solve for moments in such cases.^98–108^ Here, we use the Linear Noise Approximation (LNA) method, where assuming small fluctuations in *x_f_*(*t*) and *x_b_*(*t*) around their respective mean values 〈*x_f_*〉 and 〈*x_b_*〉, the nonlinear term is linearized as *k_b_x_f_x_b_* ≈ *k_b_*(*x_f_*〈*x_b_*〉 + 〈*x_f_*〉*x_b_* – 〈*x_b_*〉 〈*x_f_*〉). Exploiting this linearization, the probability of all events in (1) become linear with respect to the molecular counts, resulting in closed moment dynamics (see SI, section 1). A direct result of using the LNA is that the time evolution of the means is identical to the deterministic chemical rate equations

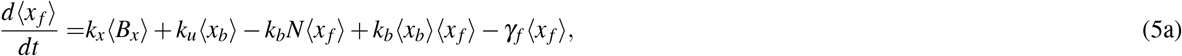

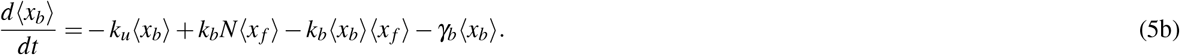

Solving the above equations at steady state, and further considering fast binding/unbinding rates (*k_b_* → ∞, *k_u_* → ∞) for a given dissociation constant *k_d_* = *k_u_*/*k_b_*, yields the following mean levels for the unbound and bound TF

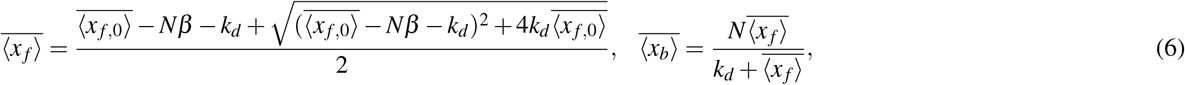

respectively. Here, 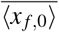 given by (3) is the mean TF count in the absence of decoy sites. When bound TFs are protected from degradation *β* = 0, the mean free TF count 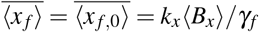 becomes independent of decoy numbers.^45,52^ In contrast, with bound TF degradation *β* > 0, the mean free TF count monotonically decreases with increasing decoy numbers, with the decrease being faster for stronger binding affinity (or lower dissociation constant). This point is exemplified in Fig. 2 where we plot 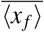 as a function of *N* for different dissociation constants. In the limit of a small number of decoys, (6) can be approximated as

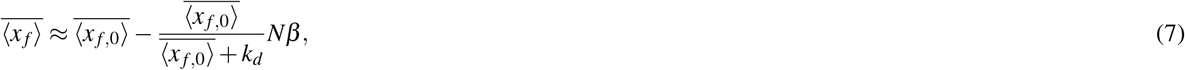

and the rate of decrease of 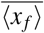 with increasing *N* is inversely proportional to *k_d_*. In the limit of large *N* and *β* > 0,

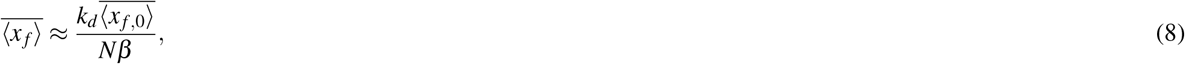

exhibiting a 1/*N* scaling of mean free TF levels with decoy abundance. Both these limits emphasize the point that when *β* > 0, increasing decoy numbers titrate away the TF, leading to reduced levels of the free TF. These results are consistent with experimental data in *Saccharomyces cerevisiae* showing reduced activity of the TF as a result of decoy binding.^37^

**Figure 2.**
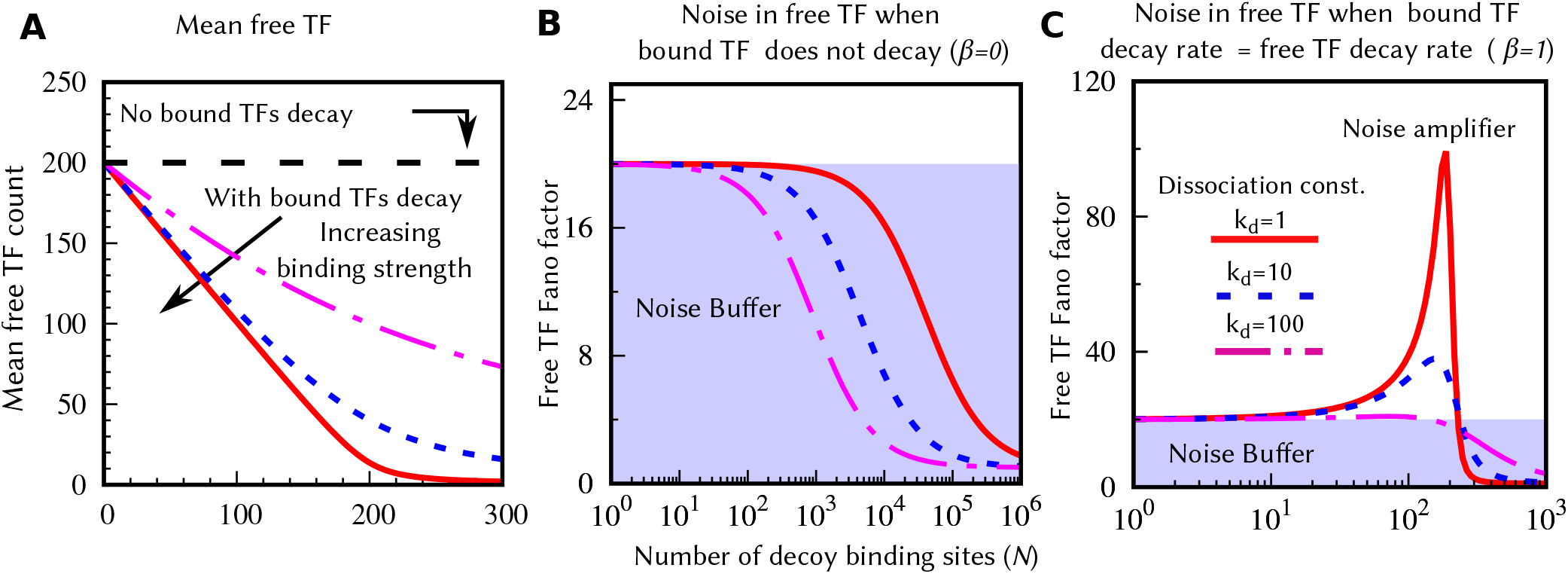
Degradation of bound TFs reduces the mean and drives non-monotonicity in the free TF noise level: The mean and the Fano factor for free TF counts are plotted against the total decoy binding sites *N* for different values of the dissociation constant (*k_d_* = 1, *k_d_* = 10 and *k_d_* = 100). (A) When bound TFs are protected from degradation, the mean becomes independent of *N* and *k_d_*. In the presence of bound TF degradation (results are shown for *β* = 1), the mean free TF count decreases with *N*. This decay becomes faster for larger binding strengths. (B) If bound TFs are protected from the degradation, the Fano factor decreases monotonically with *N*, suggesting decoys role as a noise buffer. (C) For intermediate values of decoy sites, a large noise enhancement can be seen in the presence of bound TF degradation. This noise amplification becomes larger for smaller values of *k_d_*. The Fano factors for both cases (B and C) approach to the Poissonian limit (Fano factor = 1) for large *N*. The parameters used for this figure: 〈*B_x_*〉 = 20, *k_x_* = 10 *hr*^−1^, and *γ_f_* = 1.0 *hr*^−1^ per protein molecule.

### Bound TF’s degradation enhances noise in the free TF count

Next, LNA is used to derive *F_x_f__*, the steady-state Fano factor of the free TF level in the presence of decoys. Assuming fast binding and unbinding of TFs to decoy sites (*k_b_* → ∞, *k_u_* → ∞, fixed *k_d_* = *k_u_*/*k_b_*) we obtain (see SI, section 1 for details)

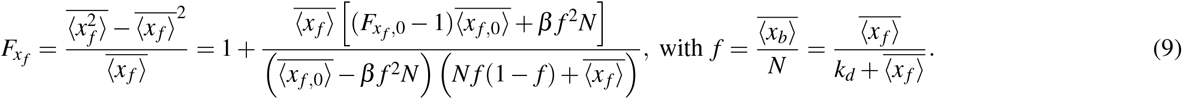

Here, *F*_*x_f_*,0_ is the Fano factor in the absence of decoy binding sites ((3)), and *f* is the fraction of bound decoys. As expected, in the limit of no decoys (*N* → 0) or weakly binding decoys (*k_d_* → ∞), *F_x_f__* → *F*_*x_f_*,0_.

When bound TFs are protected from degradation (*β* = 0), then 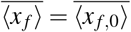 and the fraction of bound decoy *f* becomes independent of *N*. In this scenario, (9) simplifies to

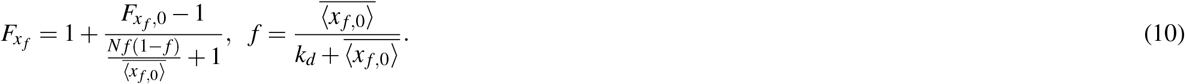

As *f* is independent of *N*, it is clear from (10) that *F_x_f__* monotonically decreases from *F*_*x_f_*,0_ to 1 with increasing *N* (Fig. 2). Thus, when the bound TFs are protected from degradation, the decoy sites function as a *noise buffer* with *F_x_f__* < *F*_*x_f_*,0_.

An interesting finding that emerges form (9) is that when *β* > 0, *F_x_f__* can vary non-monotonically with *N*. This point is illustrated in Fig. 2(C), where we plot *F_x_f__* as a function of decoy abundance for *β* = 1. While *F_x_f__* monotonically decreases to 1 for weak binding affinities (similar to the case of *β* = 0), for strong binding affinities *F_x_f__* first increases with increasing *N* to reach a maxima, before decreasing back to the Poisson limit for large *N*. In essence, with degradation of bound TFs, deocys can function as a *noise amplifier* (*F_x_f__* > *F*_*x_f_*,0_). Checking the sign of the derivative *dF_x_f__*/*dN* > 0 in the limit *N* → 0 leads to an analytical condition for noise enhancement – if the dissociation constant is below a critical threshold

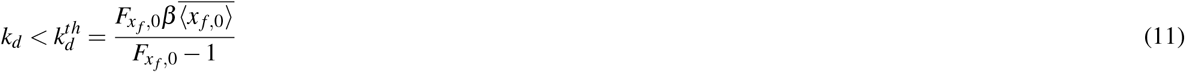

then the Fano factor will increase starting from *F*_*x_f_*,0_ as decoy sites are introduced. If *F*_*x_f_*,0_ ≫ 1, then the threshold value simplifies to 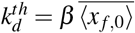 and becomes independent of *F*_*x_f_*,0_. It turns out that this condition for noise enhancement implies that the fraction of bound decoys *f* > 1/(1 + *β*) when a small number of decoys are present. In spite of the noise amplification, it is important to point out that *F_x_f__* → 1 as *N* → ∞ irrespective of the value of *β*. Thus, strongly-binding decoys that function as a noise amplifier for small *N*, become a noise buffer for large *N*.

Overall these results suggest that decoys buffer noise in a variety of scenarios: *β* = 0 (irrespective of *N* and *k_d_*), or for large number of decoys (irrespective of *β* and *k_d_*) or if decoys have sufficiently weak binding affinities (irrespective of *β* and *N*). In contrast, decoys function as a noise amplifier when *β* > 0 provided two other conditions hold – decoys have sufficiently strong binding affinities as per (11) *and* are not present in large numbers. While these observations have been made using (9) that relies on two key assumptions (LNA and fast binding/unbinding), we have confirmed the qualitative features of our results using stochastic simulations that relax both these assumptions (see SI, section 3).

As illustrated in Fig. 2, bound TFs degradation reduces the number of available TFs with increasing decoy abundance. This naturally leads to the question: is the noise behavior reported in Fig. 2 also seen when the mean free TF count is held constant at given desired level? In Fig. 3, we plot the noise as a function *N* and *k_d_* for a given mean free TF count 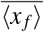 by simultaneously enhancing the production rate *k_x_* as per (6). Our results show similar qualitative behaviors with decoys functioning as a noise buffer for *β* = 0, becoming a noise amplifier when *β* = 1 for sufficiently small *N* and *k_d_* (Fig. 3). Interestingly, the region of noise amplification is greatly enhanced when bound TFs become more unstable compared to their free counterparts (Fig. 3).

**Figure 3.**
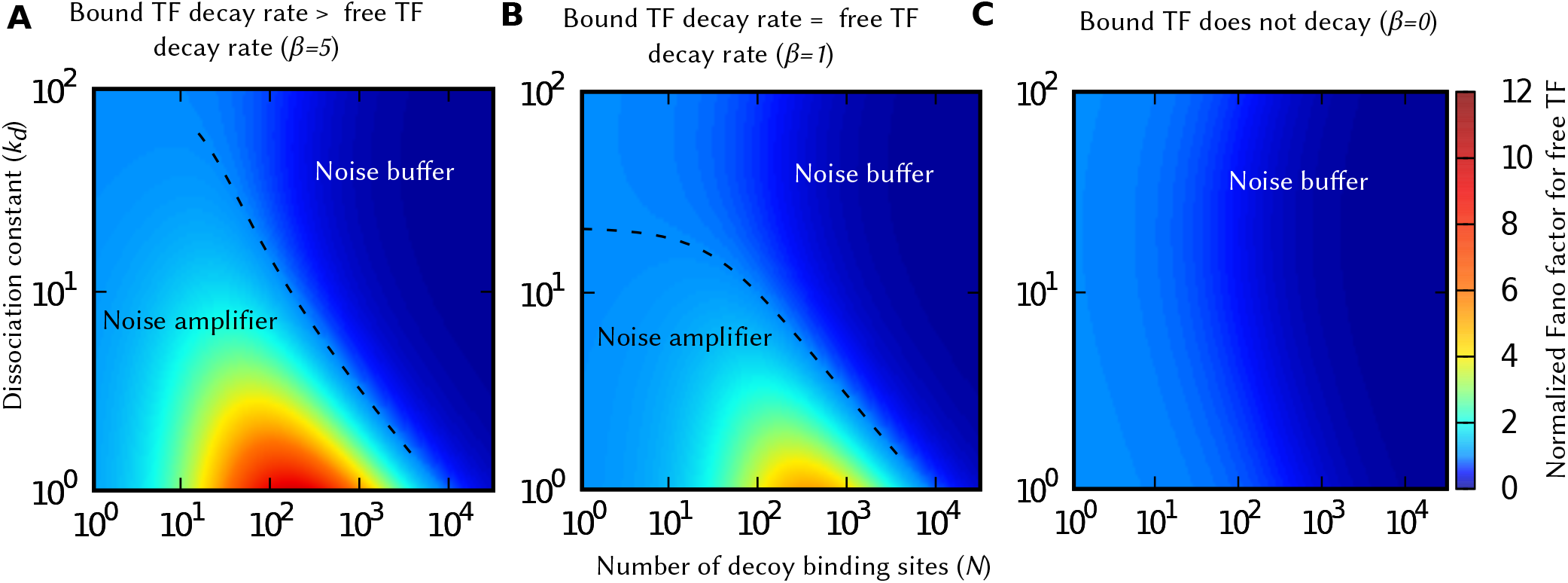
Decreasing stability of bound TFs expands the parameter space for decoy-mediated noise enhancement. The normalized Fano factors (*F_x_f__*/*F*_*x_f_*,0_) for different degradation rates of bound TFs are plotted as a function of decoy numbers (*N*) and the dissociation constant (*k_d_*), for a constant mean free TF count. The color box represents the scale for the normalized Fano factor, and its value larger than one implies noise enhancement. For smaller values of *k_d_* and *N*, the decoy acts as a noise amplifier when bound TFs are unstable. The region of noise enhancement (demarcated by a dashed line representing *F_x_f__* = *F*_*x_f_*,0_) increases with increasing degradation rate of bound TFs. Parameters used: 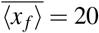 molecules, 〈*B_x_*〉 = 20, and *γ_f_* = 1 *hr*^−1^ per protein molecule. The production rate *k_x_* was varied to keep 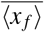 constant satisfying (6).

### Noise in free TF counts in a mixture of strong and weak decoy binding sites

Inside cells, TFs bind to various decoy sites with different affinities.^33^ How do fluctuations in the TF count gets affected by a mixture of nonidentical decoys? To address this question, we study a system two decoy types D1 and D2 that are present in numbers *N*_1_ and *N*_2_, respectively. We assume that each free TF molecule binds to both decoy types with the same (diffusion-limited) rate *k_b_*, but unbinds with different rates *k*_*u*1_ and *k*_*u*2_, respectively, leading to two different dissociation constants *k*_*d*1_ = *k*_*u*1_/*k_b_* and *k*_*d*2_ = *k*_*u*2_/*k_b_*. As before, TFs are synthesized in random bursts, the free and bound TFs decay with rates *γ_f_* and *γ_b_* (the decay rates of TFs bound to D1 and D2 are assumed to be equal). Together with (1a) and (1d), the stochastic model is now governed by the following probabilities representing jumps in the population counts in the infinitesimal time interval (*t*, *t* + *dt*]

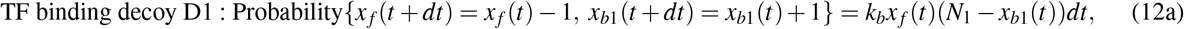

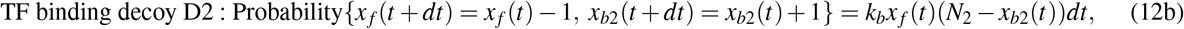

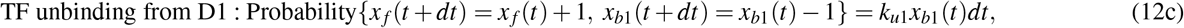

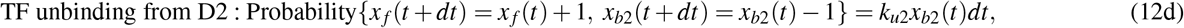

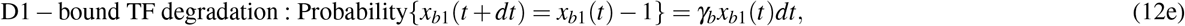

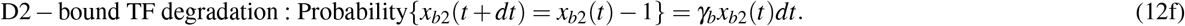

Here *x*_*b*1_ and *x*_*b*2_ denote the number TFs bound to decoys D1 and D2, respectively. We apply the linear noise approximation to obtain time evolution of statistical moments (presented in the SI, section 2), and solve the resulting moment equations numerically as the analytical formula for the noise level becomes quite involved.

Noise in the free TF counts is investigated in two complementary ways. In Fig. 4A, we fix the number of decoys D1 and their binding affinity such that D1 itself functions as a noise amplifier. The noise in the free TF counts is then plotted as a function of the number of decoys D2 and its binding affinity. Recall that the dashed line represents *F_x_f__* = *F*_*x_f_*,0_, i.e., the noise with decoys is the same as the noise in the absence of decoys. Having a large number of decoys D2 makes the overall decoy mixture a noise buffer, with the dashed line showing how large of a pool *N*_2_ is needed as a function of *k*_*d*2_. In Fig. 4B we fix the binding affinity of both decoys and plot the noise level as a function of decoys abundances. Here noise enhancement is only observed when both decoys are present at small numbers, and the decoy mixture is a noise buffer even if one of the decoys is present in sufficiently large numbers. It is interesting to note that in Fig. 4B, D1 by itself is a noise amplifier (gray line intersection with the x-axis), D2 by itself is a noise buffer (gray line intersection with the y-axis), but their combined presence (star on the dashed line) mitigates each other’s affect, and the noise level is similar to when there were no decoys.

**Figure 4.**
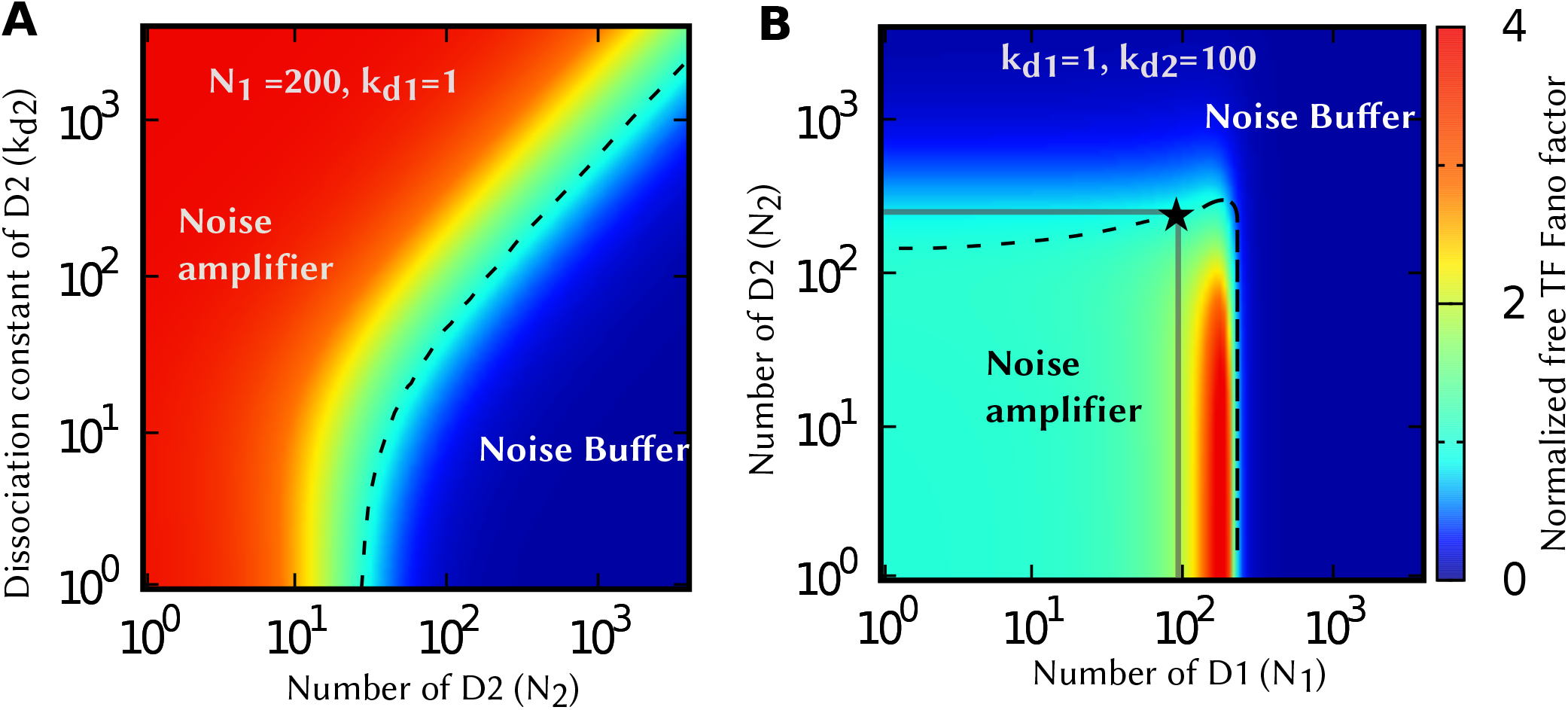
Noise-buffering decoys can mitigate the effects of noise-amplifying decoys. (A) The density plots of normalized Fano factor against *N*_2_ and *k*_*d*2_ for *N*_1_ = 200 and *k*_*d*1_ = 1. (B) The normalized Fano factor against *N*_1_ and *N*_2_ for *k*_*d*1_ = 1 and *k*_*d*2_ = 100. The noise enhancement region is separated from noise buffer region with dashed lines (correspond to *F_x_f__*/*F*_*x_f_*,0_ = 1). While individual decoy types act oppositely on free TF noise, presence of both can cancel this effect and maintain the same noise level as in the absence of any decoys. The point marked by the star is such a representative point. Parameter used: 〈*B_x_*〉 = 20, *γ_f_* = *γ_b_* = 1 *hr*^−1^ per molecule, *k_x_* = 10 *hr*^−1^, and *k_b_* = 1000 *hr*^−1^.

### Quantifying noise propagation from TF to downstream target proteins

Having quantified the magnitude of fluctuations in the free TFs counts, we next focus our attention on the time-scale of fluctuations. Given that the available TF pool activates downstream target proteins, the time scale at which fluctuations relax back to mean levels is key in understanding downstream noise propagation.^109,110^ For example, for a given noise level, more prolonged fluctuations in the free TF counts will lead to higher noise in the expression of the target protein.

The time-scale of fluctuations is quantified using the autocorrelation function defined as,

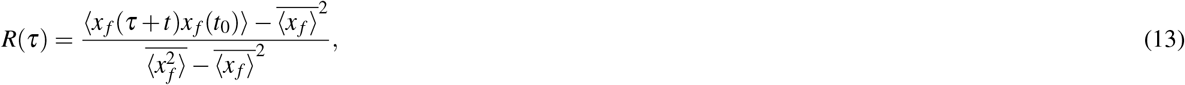

where time *t* is sufficiently large for the system to reach the steady-state. In the absence of any decoy binding sites, the autocorrelation function is simply given by exp(−*γ_f_τ*).^52^ Note that this function only depends on the decay rate and independent of bursting parameters. Thus, the magnitude and time-scale of fluctuations can be independently tuned using (3) and (13) via the decay rate, burst frequency and size. In the presence of decoy binding sites, we compute *R*(*τ*) numerically from stochastic realizations of *x_f_*(*t*) obtained via Monte Carlo simulations.^111^ Fig. 5 plots the decay of *R*(*τ*) as a function of lag-time *τ*, in the absence and presence of bound TF degradation. When the bound TFs are protected from degradations, the autocorrelation function shifts to the right leading to slower and more prolonged fluctuations in the free TF count. In contrast, when bound TFs are unstable, *R*(*τ*) shifts to the left resulting in faster fluctuations, and the shift is more pronounced for larger values of *β*.

**Figure 5.**
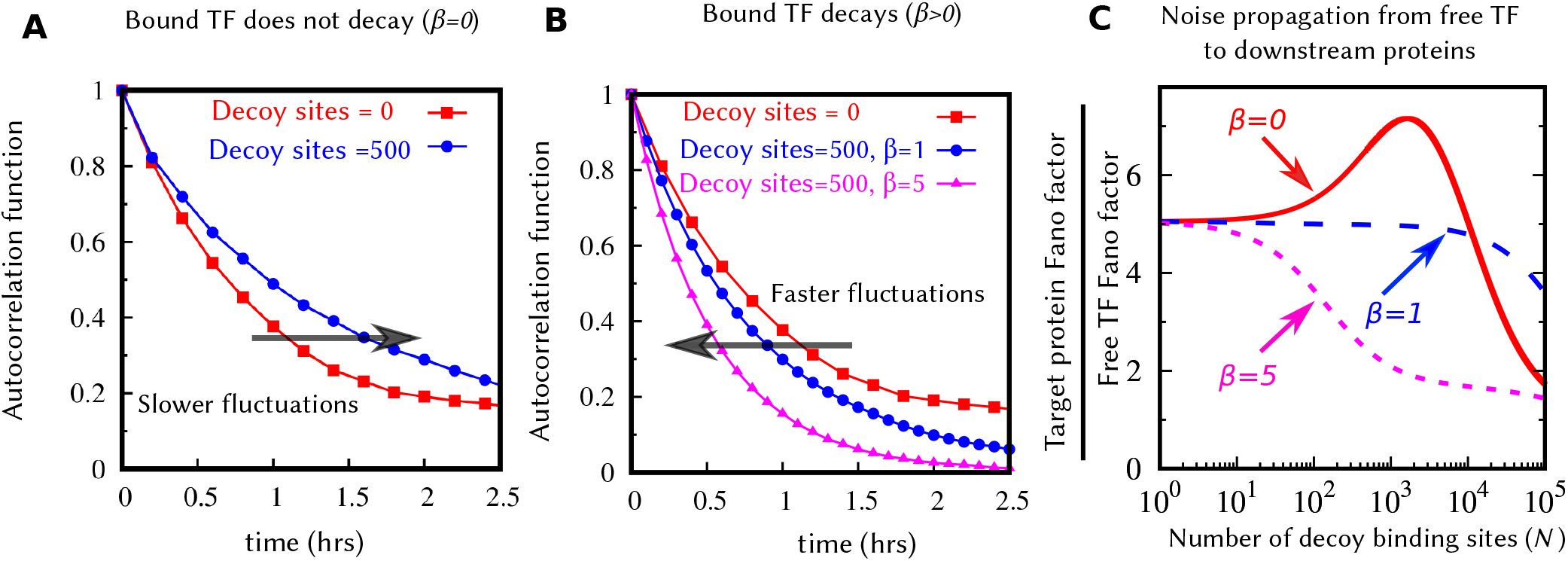
Bound TFs degradation leads to faster fluctuations in the free TF counts reducing noise propagation to downstream proteins. The autocorrelation function *R*(*τ*) of the free TF count given by (13) for (A) Bound TFs are protected from degradation, and (B) Bound TFs are subject to degradation. The addition of decoy binding sites makes the autocorrelation decay slower in (A) but faster in (B). (C) Noise propagation from the free TF to the downstream target protein is measured by the ratio of the Fano factors *F_y_*/*F_x_f__*, where *F_y_* and *F_x_f__* are given by (16) and (9), respectively. Noise propagation is enhanced when *β* = 0 consistent with the shift in the autocorrelation function to the right in Fig. 5A. In contrast, noise propagation is reduced when *β* ≥ 1. Parameters used: 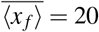 molecules, 〈*B_x_*〉 = 20, *k_d_* = 1, 〈*B_y_*〉 = 1, *k_y_* = 10*hr*^−1^, and *γ_f_* = *γ_y_* = 1*hr*^−1^ per protein molecule. We vary the *k_x_* to keep 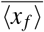 constant, satisfying (6).

To understand how fluctuations in the free TF propagate downstream, we consider stochastic synthesis of a target protein *Y* whose production is activated by available TFs via a linear dose response (see Fig. 1). The target protein is assumed to be produced in bursts *B_y_* with an arbitrary burst size distribution

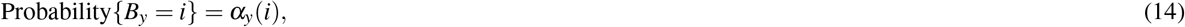

and burst frequency *x_f_k_y_* that increases linearly with the free TF count. The probabilistic events governing the production and decay of target proteins are given by

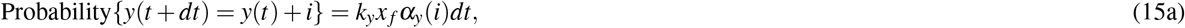

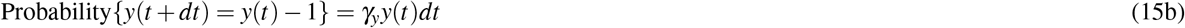

where *y*(*t*) is the population count of protein *Y* at time *t*. Applying LNA to the stochastic model (1) and (15) yields the following expression for the steady-state Fano factor of *y*(*t*) in the limit of fast binding/unbinding,

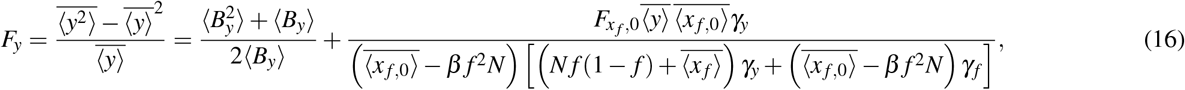

where 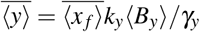 is the steady-state mean level of the target protein, 〈*B_y_*〉 is the average burst size, and 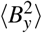 is the second-order moment of the burst size. Note that this noise level is made up of two components – the first component on the right-hand-side of (16) represents the noise from bursting of the target protein, and the second component is the noise in *Y* due to upstream fluctuations in the free TF count. In the absence of any decoy, (16) reduces to

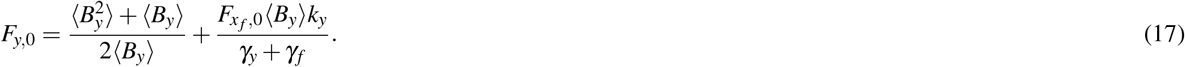

Our analysis shows that when bounds TFs are stable (*β* → 0), the second noise component in *F_y_* monotonically decreases to zero with increasing decoy numbers. When *β* > 0, then similar to the free TF count, noise enhancement in the target protein occurs for small number of high-affinity decoys, howerver both the magnitude (*F_y_*/*F*_*y*,0_) and the region of the noise enhancement are smaller compared to that of the free TF count (see Fig. S2 and compare with Fig. 3).

Noise propagation can be measured by the ratio of the target protein noise to the free TF noise, *F_y_*/*F_x_f__*. Fig. 5C shows the noise propagation as a function of decoy numbers for different decay rates of bound TF. Here the y-axis intercept quantifies noise propagation in the absence of decoys. When the bounds TFs are protected from degradation, noise propagation is first enhanced as expected from the right-shift of the autocorrelation function (Fig. 5A), but them sharply decreases at higher decoy abundances. Note that the increase in noise propagation seen at intermediate decoy numbers does not imply an increase in the target protein noise level, as at the same time the free TF noise level decreases with increasing *N* for *β* = 0 (Fig. 3). When bound TFs are unstable, noise propagation is reduced (Fig. 5C) due to faster time-scale of fluctuations of the free TF count (Fig. 5B). This implies that the noise increase seen in the free TF level when *β* > 0 (Fig. 3) is buffered by a decreased noise propagation, and hence, the region of noise enhancement for the target protein is reduced compared to that of the free TF count.

## Discussion

The sequestration of transcription factors by genomic decoy sites can either protect it from degradation^42,43^ or make it more facile for degradation.^44^ Here, we have systematically investigated how the magnitude and time-scale of TF copy-number fluctuations are impacted by the stability of the bound TF, the number and affinity of decoy sites. While this contribution focuses on TFs binding to genomic decoy sites, these results are applicable to other classes of proteins, for example RNA-binding proteins binding to sites on RNA.^112–115^

Our results show that degradation of decoy-bound TFs critically impacts both the mean and noise levels of the free TF pool. More specifically, while the average number of free (available) TFs monotonically decrease with increasing decoy abundance, the noise levels can sharply increase at low/intermediate decoy numbers before attenuating to Poisson levels as *N* → ∞ (Fig 2). This behavior can be contrasted to when bound TFs are protected from degradation, in which case the mean free TF count becomes invariant of *N* and decoys always buffer noise (Figs. 2 and 3). When *β* > 0, high-affinity decoys can transition from being noise amplifiers to noise buffers as their numbers are increased (Fig. 2). This point is exemplified in Fig. 6 where high-affinity low-abundance decoys, and low-affinity high-abundance decoys result in the same average number of freely available TFs, but with much higher noise in the former case. Moreover, a mixture of both decoy types can mitigate each other’s affect leading to no noise enhancement or buffering (Fig. 4).

**Figure 6.**
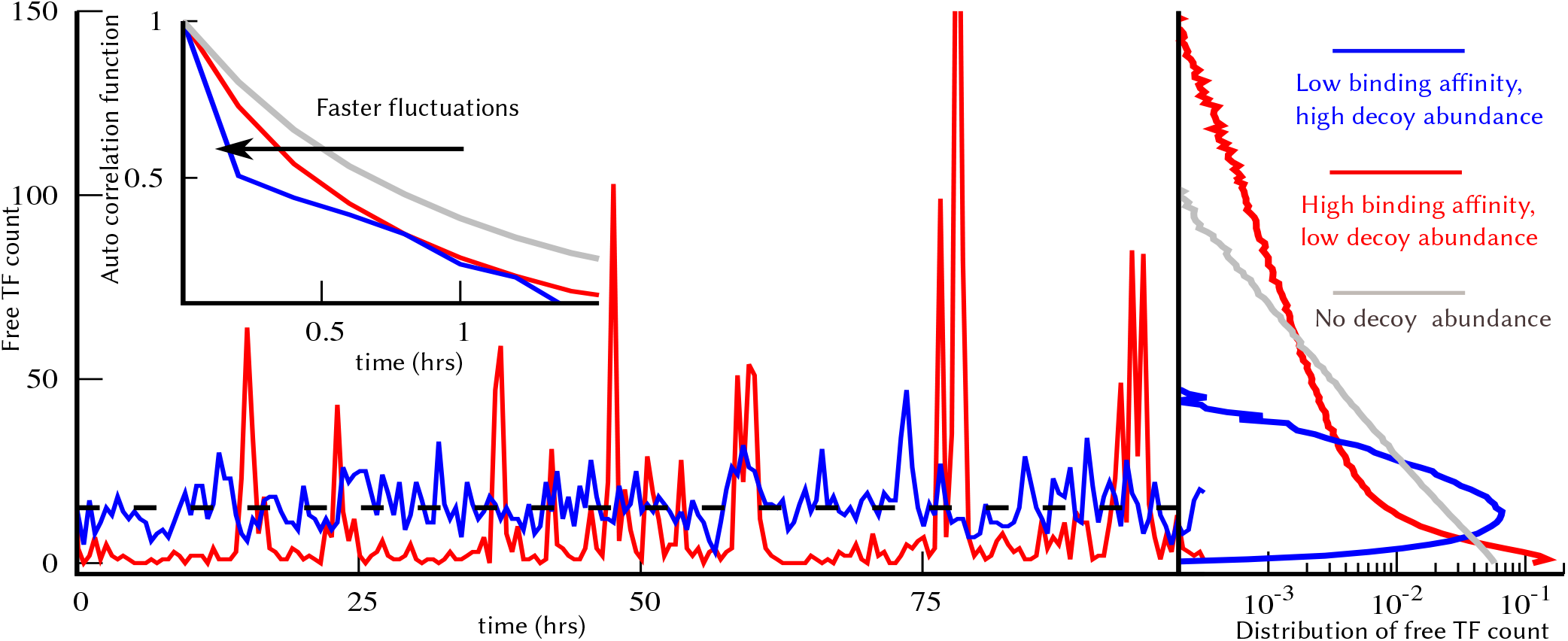
High-affinity low-abundance decoys, and low-affinity high-abundance decoys result in the same mean free TF counts with contrasting fluctuation dynamics. Stochastic realizations of the number of free TFs for high-affinity low-abundance decoys (red), and low-affinity high-abundance decoys (blue) when *β* = 1, along with their corresponding probability distributions on the right. Both scenarios yield the same average number of free TF molecules, but high-affinity decoys drive significantly enhanced noise levels. The instability of the bound TF leads to faster time-scale of fluctuations as illustrated by the left-shift of the autocorrelation function (inset). For comparison, the free TF count distribution and the autocorrelation function for the no-decoy case are also shown with grey lines. The parameters used are as follows: 〈*B_x_*〉 = 20, *k_x_* = 10 *hr*^−1^, *γ_f_* = *γ_b_* = 1.0 *hr*^−1^ per protein molecule. For high affinity decoy *k_d_* = 1 and *N* = 245, and low affinity decoy *k_d_* = 100 and *N* = 1400. This choice produce the same mean TF count for both the cases, 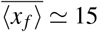. For the no-decoy case: *k_x_* = 0.75*hr*^−1^ to keep 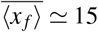.

Closed-form formulas for the noise levels derived using the Linear Noise Approximation led to precise conditions for noise enhancement – the number of decoys is not large, and their binding affinity is strong enough such that the fraction of bound decoys is higher than 1/(*β* + 1). For example, *β* = 1 for a stable TF whoes turnover is primarily governed by dilution from cell growth, and this condition implies that more than 50% decoy sites must be occupied. The decoy-mediated noise enhancement is higher for large values of *β*, and for more “sticky” or higher affinity decoys (Fig. 3). It is interesting to point out that the peak noise enhancement generally occurs when the number of decoy sites is comparable to the TF counts. For example, in Fig. 2C there are an average of 200 TF molecules in the absence of decoys and the noise peak is also seen around *N* ≈ 200. Note that our results are restricted to TF production in stochastic bursts, and it remains to be seen if these results generalize to other noise mechanisms in gene expression, such as, extrinsic noise that arises from intercellular differences in cell size and the abundance of expression machinery.^116–120^

The speed of fluctuation in free TF counts also shows distinct features in the presence and absence of bound TF degradation. While the fluctuations become faster in the presence of bound TF degradation, as indicated by the left-shift of the autocorrelation function (Fig. 5), we see an opposite result when bound TFs do not degrade. These results have important implications on how noise propagates from the TF to downstream target proteins. For example, when *β* > 0 decoys can amplify noise in the free TF levels, while at the same time make fluctuations in TF counts relax back faster attenuating downstream noise transmission. As a result of these two opposing forces, the noise in the target protein may not increase even though the noise in the free TF level has increased. This can be seen for *β* = 5 in Fig. 3 where for *N* = 100 and *k_d_* = 10 the decoy is a noise amplifier, but as seen in Fig. S2, for the exact same parameter values the noise in the target protein has reduced compared to the no-decoy case.

Our novel finding of the dual role of decoys as noise amplifiers/buffers clearly encourages more investigation into the regulatory function of decoys in complex gene networks. For example, in genetic/signalling circuits with oscillatory dynamics decoys can tune the oscillation frequency^48^, enhance coherence,^47^ and it will be interesting to see if decoys can make biological clocks more robust to molecular noise. An exciting avenue of experimental research is to use decoys as manipulations of phenotypic heterogeneity. For example, aberrant expression of resistance markers in individual melanoma cells have been shown to be drives of cancer drug resistance,^121^ and decoy-mediated noise buffering can play a therapeutic role in reducing such outlier cells. In the context of the Human Immunodeficiency Virus (HIV), noisy expression of a key viral protein, Tat, controls the cell-fate outcome between active viral replication and latency.^16,122–125^ Latency is a dormant state of the virus and considered the biggest challenge in eradicating the virus from a patient since latent cells are long lived and resistant to drug therapy.^126^ Recently, small-molecule compounds have been identified that enhance noise in Tat expression for efficient reactivation of latent cells,^127^ and here the role of decoys as noise amplifiers allows for a Tat-specific compound-free approach for latency reversal.

## Supplementary Information

### 1. Dynamical equations for moments for free and bound TFs and target proteins

Let *ϕ*(*x_f_*, *x_b_*) be an arbitrary differentiable function of free TF count *x_f_*, bound TF count *x_b_*. The time evolution of *ϕ*(*x_f_*, *x_b_*) obeying the chemical master equation Eq. (2) in the main text is given by,

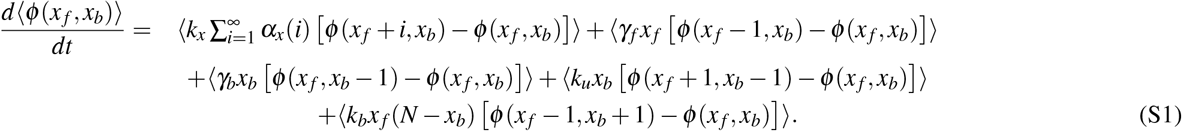

Note that the last term (due to binding event) is nonlinear. This makes the moment dynamics unclosed in the sense that the dynamical equation of a given moment depends on the higher order moments. For example, the time evolution of 〈*x_f_*〉, which can be obtained by substituting *ϕ*(*x_f_*, *x_b_*) with *x_f_* in (S1), is given by,

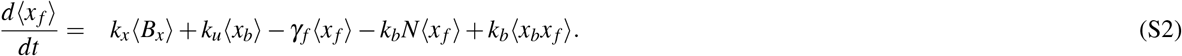

This dynamical equation of first order moment depends higher order moment 〈*x_b_x_f_*〉. Similarly, the dynamical equation of 〈*x_b_x_f_*〉 depends on the higher order moments such as 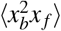 and 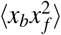. To get closed form moment equations, we use the Linear Noise Approximation (LNA). Under this approximation, we linearise the nonlinear part of the binding term *k_b_x_b_x_f_* around the mean levels 〈*x_f_*〉 and 〈*x_b_*〉: *k_b_x_b_x_f_* = *k_b_*(〈*x_b_*〉*x_f_* + *x_b_*〈*x_f_*〉 – 〈*x_b_*〉〈*x_f_*〉). Under the LNA, the first order moment dynamics of *x_f_* and *x_b_* are given by,

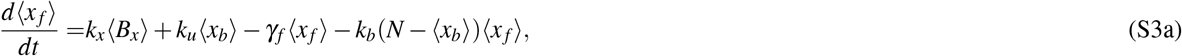

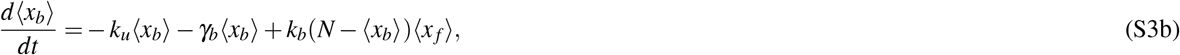

The steady value of the mean free TF and bound TF are obtained by solving the above equation with setting time derivative values to zero. In the limit of large binding/unbinding rates (*k_b_* → ∞, *k_u_* → ∞ with *k_u_*/*k_b_* = *k_d_*), the expression of 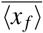, and 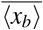 are presented in the main text. We use the same linearization scheme to obtain the dynamics for second moments,

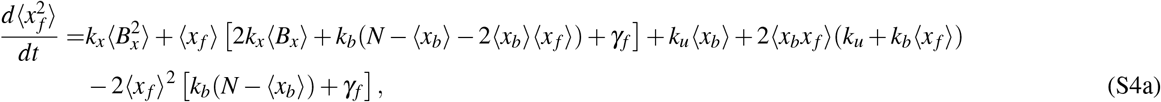

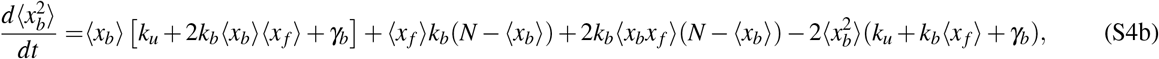

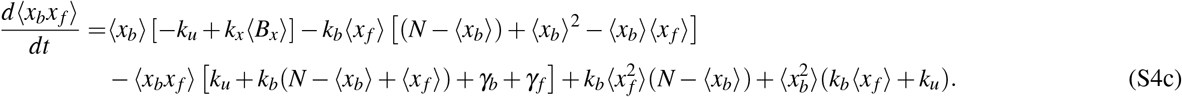

In the case of downstream protein (*Y*), the additional first and second order moment dynamics are given by,

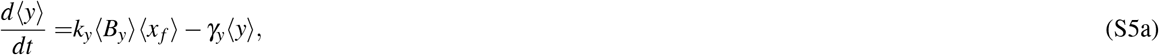

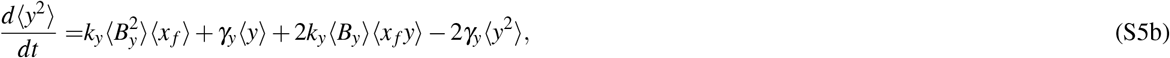

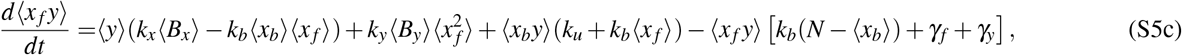

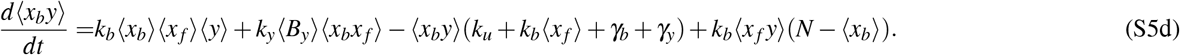

We solve (S3),(S4) and (S5) at the steady state, and in the limit of large binding/unbinding rates, we derive the expressions the Fano factors for free TF count and target protein count which are presented in the main text.

### 2. Dynamical equations for two decoy species

In the case of two decoy species, the dynamical equations of the first order moments are given by,

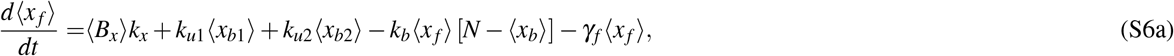

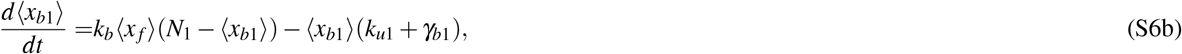

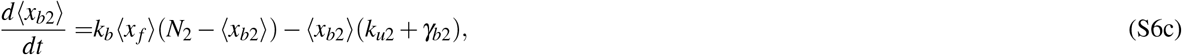

where *N* = *N*_1_ + *N*_2_ is the decoy sites, and *x_b_* = *x*_*b*1_ + *x*_*b*2_ is the total bound complexes. The dynamical equations for the second order moments are

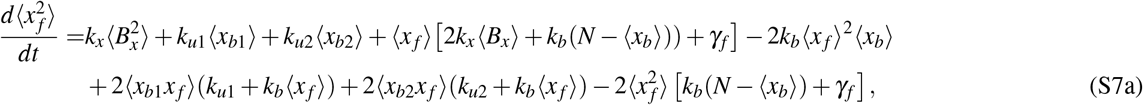

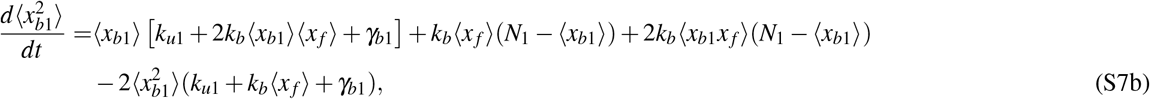

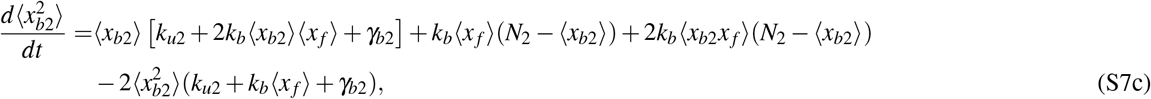

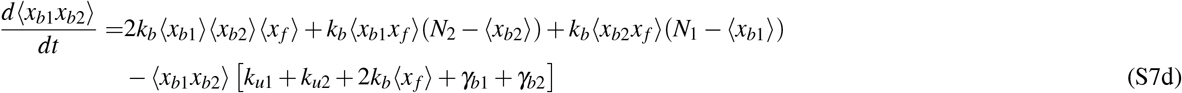

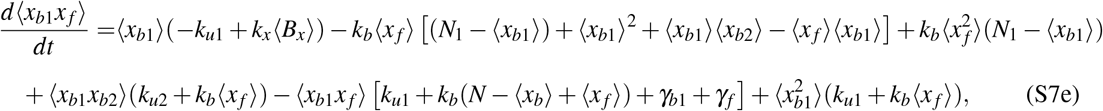

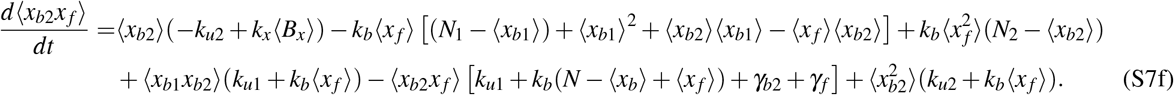

In the steady state, for a given set of parameters, we solve these equations numerically to obtain the density plot presented in the main text.

### 3. The comparison between LNA and exact stochastic simulation results

We use kinetic Monte Carlo algorithm due to Gillespie [Gillespie, Daniel T, *Journal of Comput. Phys*. **22**, 403–434 (1976)] to solve our stochastic model numerically. For a large dissociation constant, the LNA results for the free TF Fano factor match with the simulation results quite well. However, for a small dissociation constant, a clear deviation is observed near the peak position. However, the qualitative behavior is similar.

**Figure S1.**
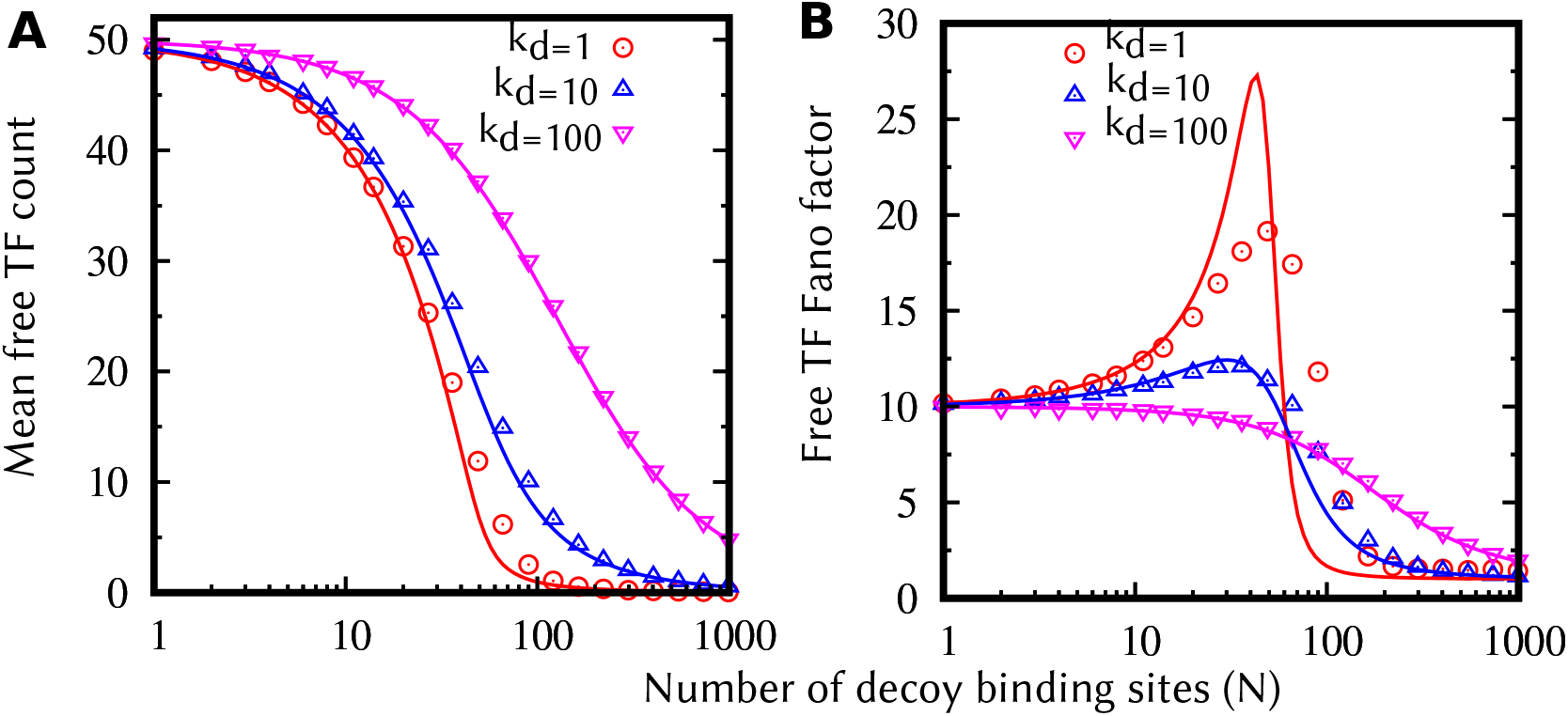
The comparison of the LNA and stochastic simulation results: The mean free TF counts (A) and the Fano factor (B) as a function of the total number of decoy binding sites for three values of binding affinities. The data points represented by symbols are obtained from stochastic simulations and the corresponding lines plotted using the Fano factor formula for free TF presented in the main text. Parameters used: *k_x_* = 5 *hr*^−1^ molecules, 〈*B_x_*〉 = 10, and *γ_b_* = *γ_f_* = 1 *hr*^−1^ per protein molecule.

### 4. Density plots for noise in the traget protein

**Figure S2.**
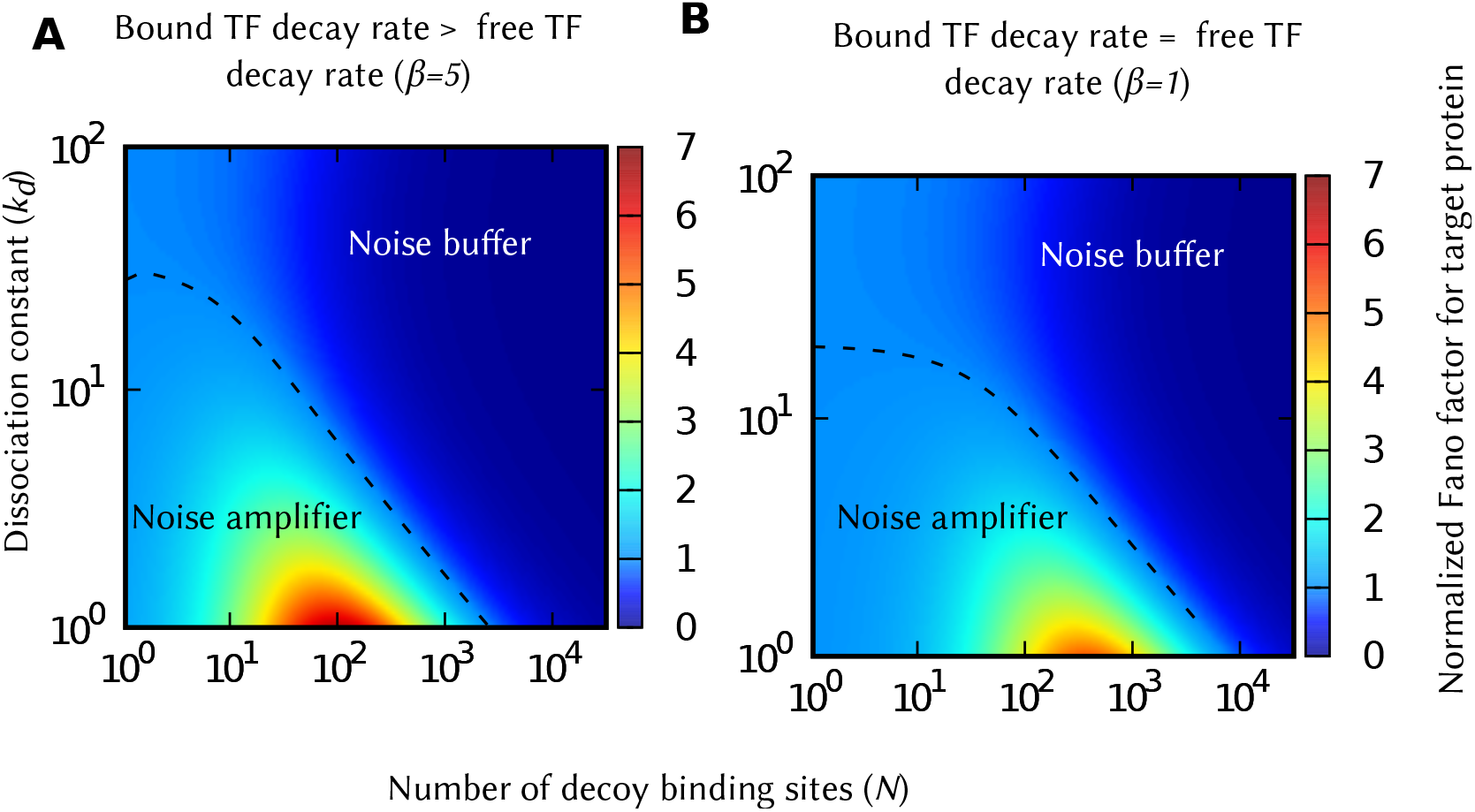
The normalized Fano factor (*F_y_*/*F*_*y*,0_) for the target protein for constant values 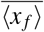 and 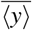. The target protein counts show noise enhancement similar to free TF count. However, the magnitude and the region of the noise enhancement are relatively smaller to that of free TF count. Parameters used: 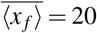 and 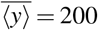 molecules, 〈*B_x_*〉 = 20, and *γ_y_* = *γ_f_* = 1 *hr*^−1^ per protein molecule. We vary the *k_x_* to keep 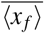 constant satisfying (6).

## Acknowledgments

AS is supported by the NSF grant ECCS-1711548, and NIH grants 5R01GM124446 and 5R01GM126557.

## References

1. McAdams, H. H. & Arkin, A. It’s a noisy business! Genetic regulation at the nanomolar scale. Trends in genetics 15, 65–9 (1999).

2. Arkin, A. P., Ross, J. & McAdams, H. H. Stochastic kinetic analysis of developmental pathway bifurcation in phage λ-infected *Escherichia coli* cells. Genetics 149, 1633–1648 (1998).

3. Elowitz, M. B., Levine, A. J., Siggia, E. D. & Swain, P. S. Stochastic gene expression in a single cell. Science 297, 1183–1186 (2002).

4. Losick, R. & Desplan, C. Stochasticity and cell fate. Science 320, 65–68 (2008).

5. Shaffer, S. M. et al. Rare cell variability and drug-induced reprogramming as a mode of cancer drug resistance. Nature 546, 431–435 (2017).

6. Billman, M., Rueda, D. & Bangham, C. Single-cell heterogeneity and cell-cycle-related viral gene bursts in the human leukaemia virus htlv-1. Wellcome Open Research 2, 87 (2017).

7. Keskin, S. et al. Noise in the vertebrate segmentation clock is boosted by time delays but tamed by notch signaling. Cell Rep 23, 2175–2185 (2018).

8. Urban, E. A. & Johnston, R. J. Buffering and amplifying transcriptional noise during cell fate specification. Frontiers in Genetics 9, 591 (2018).

9. Raj, A. & van Oudenaarden, A. Nature, nurture, or chance: stochastic gene expression and its consequences. Cell 135, 216–226 (2008).

10. von Dassow, G., Meir, E., Munro, E. M. & Odell, G. M. The segment polarity network is a robust developmental module. Nature 406, 188 (2000).

11. Queitsch, C., Sangster, T. A. & Lindquist, S. Hsp90 as a capacitor of phenotypic variation. Nature 417, 618 (2002).

12. Schmiedel, J. M., Carey, L. B. & Lehner, B. Empirical mean-noise fitness landscapes reveal the fitness impact of gene expression noise. Nature Communications 10, 3180 (2019).

13. Kemkemer, R., Schrank, S., Vogel, W., Gruler, H. & Kaufmann, D. Increased noise as an effect of haploinsufficiency of the tumor-suppressor gene neurofibromatosis type 1 in vitro. Proceedings of the National Academy of Sciences 99, 13783–13788 (2002).

14. Bahar, R. et al. Increased cell-to-cell variation in gene expression in ageing mouse heart. Nature 441, 1011–1014 (2006).

15. Kussell, E. & Leibler, S. Phenotypic diversity, population growth, and information in fluctuating environments. Science 309, 2075–2078 (2005).

16. Weinberger, L. S., Burnett, J., Toettcher, J., Arkin, A. & Schaffer, D. Stochastic gene expression in a lentiviral positive-feedback loop: HIV-1 Tat fluctuations drive phenotypic diversity. Cell 122, 169–182 (2005).

17. Kærn, M., Elston, T. C., Blake, W. J. & Collins, J. J. Stochasticity in gene expression: from theories to phenotypes. Nature Reviews Genetics 6, 451–464 (2005).

18. Blake, W. J. et al. Phenotypic consequences of promoter-mediated transcriptional noise. Molecular Cell 24, 853–865 (2006).

19. Davidson, C. J. & Surette, M. G. Individuality in bacteria. Annual Review of Genetics 42, 253–268 (2008).

20. Fraser, D. & Kaern, M. A chance at survival: gene expression noise and phenotypic diversification strategies. Mol Microbiol 71, 1333–1340 (2009).

21. Rotem, E. et al. Regulation of phenotypic variability by a threshold-based mechanism underlies bacterial persistence. Proceedings of the National Academy of Sciences 107, 12541–12546 (2010).

22. Schreiber, F. et al. Phenotypic heterogeneity driven by nutrient limitation promotes growth in fluctuating environments. Nature Microbiology 1, 16055 (2016).

23. Maamar, H., Raj, A. & Dubnau, D. Noise in gene expression determines cell fate in bacillus subtilis. Science 317, 526–529 (2007).

24. Balázsi, G., van Oudenaarden, A. & Collins, J. J. Cellular decision making and biological noise: From microbes to mammals. Cell 144, 910–925 (2014).

25. Ozbudak, E. M., Thattai, M., Kurtser, I., Grossman, A. D. & van Oudenaarden, A. Regulation of noise in the expression of a single gene. Nature Genetics 31, 69–73 (2002).

26. Singh, A. & Hespanha, J. P. Optimal feedback strength for noise suppression in autoregulatory gene networks. Biophysical Journal 96, 4013–4023 (2009).

27. Dublanche, Y., Michalodimitrakis, K., Kummerer, N., Foglierini, M. & Serrano, L. Noise in transcription negative feedback loops: simulation and experimental analysis. Molecular Systems Biology 2, 41 (2006).

28. Blake, W. J., Kaern, M., Cantor, C. R. & Collins, J. J. Noise in eukaryotic gene expression. Nature 422, 633–637 (2003).

29. Alon, U. An Introduction to Systems Biology: Design Principles of Biological Circuits (Chapman and Hall/CRC, 2011).

30. Bintu, L. et al. Transcriptional regulation by the numbers: applications. Current Opinion in Genetics & Development 15, 125 – 135 (2005).

31. Sánchez, Á. & Kondev, J. Transcriptional control of noise in gene expression. Proceedings of the National Academy of Sciences 105, 5081–5086 (2008).

32. Wunderlich, Z. & Mirny, L. A. Different gene regulation strategies revealed by analysis of binding motifs. Trends in genetics 25, 434–440 (2009).

33. Kemme, C. A., Nguyen, D., Chattopadhyay, A. & Iwahara, J. Regulation of transcription factors via natural decoys in genomic dna. Transcription 7, 115–120 (2016).

34. Esadze, A., Kemme, C. A., Kolomeisky, A. B. & Iwahara, J. Positive and negative impacts of nonspecific sites during target location by a sequence-specific dna-binding protein: origin of the optimal search at physiological ionic strength. Nucleic Acids Research 42, 7039–7046 (2014).

35. Kemme, C. A., Esadze, A. & Iwahara, J. Influence of quasi-specific sites on kinetics of target dna search by a sequencespecific dna-binding protein. Biochemistry 54, 6684–6691 (2015).

36. Bakk, A. & Metzler, R. In vivo non-specific binding of λ ci and cro repressors is significant. FEBS Letters 563, 66 – 68 (2004).

37. Lee, T. & Maheshri, N. A regulatory role for repeated decoy transcription factor binding sites in target gene expression. Molecular systems biology 8, 576 (2012).

38. Morishita, R. et al. A gene therapy strategy using a transcription factor decoy of the e2f binding site inhibits smooth muscle proliferation in vivo. Proceedings of the National Academy of Sciences 92, 5855–5859 (1995).

39. Mann, M. J. Transcription factor decoys: A new model for disease intervention. Annals of the New York Academy of Sciences 1058, 128–139 (2005).

40. Hecker, M. & Wagner, A. H. Transcription factor decoy technology: A therapeutic update. Biochemical Pharmacology 144, 29 – 34 (2017).

41. Francois, M., Donovan, P. & Fontaine, F. Modulating transcription factor activity: Interfering with protein-protein interaction networks. Seminars in Cell and Developmental Biology (2018).

42. Abu Hatoum, O. et al. Degradation of myogenic transcription factor myod by the ubiquitin pathway in vivo and in vitro: Regulation by specific dna binding. Molecular and Cellular Biology 18, 5670–5677 (1998).

43. Molinari, E., Gilman, M. & Natesan, S. Proteasome-mediated degradation of transcriptional activators correlates with activation domain potency in vivo. EMBO J 18, 6439–6447 (1999).

44. Thomas, D. & Tyers, M. Transcriptional regulation: Kamikaze activators. Current Biology 10, R341 – R343 (2000).

45. Burger, A., Walczak, A. M. & Wolynes, P. G. Abduction and asylum in the lives of transcription factors. Proceedings of the National Academy of Sciences 107, 4016–4021 (2010).

46. Burger, A., Walczak, A. M. & Wolynes, P. G. Influence of decoys on the noise and dynamics of gene expression. Physical Review E 86, 041920 (2012).

47. Wang, Z., Potoyan, D. A. & Wolynes, P. G. Molecular stripping, targets and decoys as modulators of oscillations in the nf-kb/ikbα/dna genetic network. Journal of The Royal Society Interface 13, 20160606 (2016).

48. Jayanthi, S. & Del Vecchio, D. Tuning genetic clocks employing DNA binding sites. PLOS ONE 7, e41019 (2012).

49. Jayanthi, S., Nilgiriwala, K. S. & Del Vecchio, D. Retroactivity controls the temporal dynamics of gene transcription. ACS Synthetic Biology 2, 431–441 (2013).

50. Ricci, F., Vallée-Bélisle, A. & Plaxco, K. W. High-precision, in vitro validation of the sequestration mechanism for generating ultrasensitive dose-response curves in regulatory networks. PLOS Computational Biology 7, e1002171 (2011).

51. Jones, D. L., Brewster, R. C. & Phillips, R. Promoter architecture dictates cell-to-cell variability in gene expression. Science 346, 1533–1537 (2014).

52. Soltani, M., Bokes, P., Fox, Z. & Singh, A. Nonspecific transcription factor binding can reduce noise in the expression of downstream proteins. Physical Biology 12, 055002 (2015).

53. Bokes, P. & Singh, A. Protein copy number distributions for a self-regulating gene in the presence of decoy binding sites. PLOS ONE 10, e0120555 (2015).

54. Das, D., Dey, S., Brewster, R. C. & Choube, S. Effect of transcription factor resource sharing on gene expression noise. PLoS Comput Biol 13, e1005491 (2017).

55. Razo-Mejia, M. et al. Tuning transcriptional regulation through signaling: A predictive theory of allosteric induction. Cell Systems 6, 456 – 469 (2018).

56. Suter, D. M. et al. Mammalian genes are transcribed with widely different bursting kinetics. Science 332, 472–474 (2011).

57. Dar, R. D. et al. Transcriptional burst frequency and burst size are equally modulated across the human genome. Proceedings of the National Academy of Sciences 109, 17454–17459 (2012).

58. Fukaya, T., Lim, B. & Levine, M. Enhancer control of transcriptional bursting. Cell 166, 358–368 (2015).

59. Bartman, C. R., Hsu, S. C., Hsiung, C. C.-S., Raj, A. & Blobel, G. A. Enhancer regulation of transcriptional bursting parameters revealed by forced chromatin looping. Molecular Cell 62, 237 – 247 (2016).

60. Corrigan, A. M., Tunnacliffe, E., Cannon, D. & Chubb, J. R. A continuum model of transcriptional bursting. eLife 5, e13051 (2016).

61. Chong, S., Chen, C., Ge, H. & Xie, X. S. Mechanism of transcriptional bursting in bacteria. Cell 158, 314–326 (2014).

62. Singh, A., Razooky, B., Cox, C. D., Simpson, M. L. & Weinberger, L. S. Transcriptional bursting from the HIV-1 promoter is a significant source of stochastic noise in HIV-1 gene expression. Biophysical Journal 98, L32–L34 (2010).

63. Dar, R. D. et al. Transcriptional bursting explains the noise–versus–mean relationship in mRNA and protein levels. PLOS ONE 11, e0158298 (2016).

64. Golding, I., Paulsson, J., Zawilski, S. & Cox, E. Real-time kinetics of gene activity in individual bacteria. Cell 123, 1025–1036 (2005).

65. Raj, A., Peskin, C., Tranchina, D., Vargas, D. & Tyagi, S. Stochastic mRNA synthesis in mammalian cells. PLOS Biology 4, e309 (2006).

66. Singh, A., Razooky, B. S., Dar, R. D. & Weinberger, L. S. Dynamics of protein noise can distinguish between alternate sources of gene-expression variability. Molecular Systems Biology 8, 607 (2012).

67. Thattai, M. & van Oudenaarden, A. Intrinsic noise in gene regulatory networks. Proceedings of the National Academy of Sciences 98, 8614–8619 (2001).

68. Friedman, N., Cai, L. & Xie, X. Linking stochastic dynamics to population distribution: an analytical framework of gene expression. Physical Review Letters 97, 168302 (2006).

69. Shahrezaei, V. & Swain, P. S. Analytical distributions for stochastic gene expression. Proceedings of the National Academy of Sciences 105, 17256–17261 (2008).

70. Pedraza, J. M. & Paulsson, J. Effects of molecular memory and bursting on fluctuations in gene expression. Science 319, 339 – 343 (2008).

71. Jia, T. & Kulkarni, R. V. Intrinsic noise in stochastic models of gene expression with molecular memory and bursting. Journal of Mathematical Biology 106, 058102 (2011).

72. Kumar, N., Singh, A. & Kulkarni, R. V. Transcriptional bursting in gene expression: Analytical results for genera stochastic models. PLOS Computational Biology 11, e1004292 (2015).

73. Bokes, P. & Singh, A. Gene expression noise is affected deferentially by feedback in burst frequency and burst size. Journal of Mathematical Biology 74, 1483–1509 (2017).

74. Singh, A. & Soltani, M. Quantifying intrinsic and extrinsic variability in stochastic gene expression models. PLOS ONE 8, e84301 (2013).

75. Soltani, M., Vargas-Garcia, C. A., Antunes, D. & Singh, A. Intercellular variability in protein levels from stochastic expression and noisy cell cycle processes. PLOS Computational Biology e1004972 (2016).

76. Yu, J., Xiao, J., Ren, X., Lao, K. & Xie, X. S. Probing gene expression in live cells, one protein molecule at a time. Science 311, 1600–1603 (2006).

77. Paulsson, J. Model of stochastic gene expression. Physics of Life Reviews 2, 157–175 (2005).

78. Elgart, V., Jia, T., Fenley, A. T. & Kulkarni, R. Connecting protein and mrna burst distributions for stochastic models of gene expression. Physical biology 8, 046001 (2011).

79. Wilkinson, D. J. Stochastic Modelling for Systems Biology (Chapman and Hall/CRC, 2011).

80. McQuarrie, D. A. Stochastic approach to chemical kinetics. Journal of Applied Probability 4, 413–478 (1967).

81. Munsky, B. & Khammash, M. The finite state projection algorithm for the solution of the chemical master equation. Journal of Chemical Physics 124, 044104 (2006).

82. Gupta, A., Mikelson, J. & Khammash, M. A finite state projection algorithm for the stationary solution of the chemical master equation. The Journal of Chemical Physics 147, 154101 (2017).

83. Dinh, K. N. & Sidje, R. B. Understanding the finite state projection and related methods for solving the chemical master equation. Physical Biology 13, 035003 (2016).

84. Gillespie, D. T. Approximate accelerated stochastic simulation of chemically reacting systems. Journal of Chemical Physics 115, 1716–1733 (2001).

85. Gibson, M. A. & Bruck, J. Efficient exact stochastic simulation of chemical systems with many species and many channels. Journal of Physical Chemistry A 104, 1876–1889 (2000).

86. Cao, Y., Li, H. & Petzold, L. Efficient formulation of the stochastic simulation algorithm for chemically reacting systems. Journal of Chemical Physics 121, 4059–4067 (2004).

87. Anderson, D. F. A modified next reaction method for simulating chemical systems with time dependent propensities and delays. Journal of Chemical Physics 127, 214107 (2007).

88. Daigle, B., Soltani, M., Petzold, L. & Singh, A. Inferring single-cell gene expression mechanisms using stochastic simulation. Bioinformatics 31, 1428–1435 (2015).

89. Van Kampen, N. Stochastic Processes in Physics and Chemistry (Elsevier, 2011).

90. Elf, J. & Ehrenberg, M. Fast evaluation of fluctuations in biochemical networks with the linear noise approximation. Genome Research 13, 2475–2484 (2003).

91. Munsky, B., Hlavacek, W. S. & Tsimring, L. S. Quantitative biology: theory, computational methods, and models (The MIT Press, 2018).

92. Modi, S., Soltani, M. & Singh, A. Linear noise approximation for a class of piecewise deterministic markov processes. In 2018 Annual American Control Conference (ACC), 1993–1998 (2018).

93. Elf, J. & Ehrenberg, M. Fast evaluation of fluctuations in biochemical networks with the linear noise approximation. Genome Research 13, 2475–2484 (2003).

94. Thomas, P., Straube, A. V. & Grima, R. The slow-scale linear noise approximation: an accurate, reduced stochastic description of biochemical networks under timescale separation conditions. BMC Systems Biology 6, 39 (2012).

95. Singh, A. Transient changes in intercellular protein variability identify sources of noise in gene expression. Biophysical Journal 107, 2214–2220 (2014).

96. Singh, A. & Hespanha, J. P. Stochastic hybrid systems for studying biochemical processes. Philosophical Transactions of the Royal Society A 368, 4995–5011 (2010).

97. Singh, A. & Hespanha, J. P. Approximate moment dynamics for chemically reacting systems. IEEE Transactions on Automatic Control 56, 414–418 (2011).

98. Gomez-Uribe, C. A. & Verghese, G. C. Mass fluctuation kinetics: Capturing stochastic effects in systems of chemical reactions through coupled mean-variance computations. Journal of Chemical Physics 126, 024109 (2007).

99. Lee, C. H., Kim, K. & Kim, P. A moment closure method for stochastic reaction networks. Journal of Chemical Physics 130, 134107 (2009).

100. Goutsias, J. Classical versus stochastic kinetics modeling of biochemical reaction systems. Biophysical Journal 92, 2350–2365 (2007).

101. Gillespie, C. S. Moment closure approximations for mass-action models. IET Systems Biology 3, 52–58 (2009).

102. Soltani, M., Vargas, C. & Singh, A. Conditional moment closure schemes for studying stochastic dynamics of genetic circuits. IEEE Transactions on Biomedical Systems and Circuits 9, 518–526 (2015).

103. Zhang, J., DeVille, L., Dhople, S. & Dominguez-Garcia, A. A maximum entropy approach to the moment closure problem for stochastic hybrid systems at equilibrium. In IEEE Conference on Decision and Control, 747–752 (2014).

104. Smadbeck, P. & Kaznessis, Y. N. A closure scheme for chemical master equations. Proceedings of the National Academy of Sciences 110, 14261–14265 (2013).

105. Schnoerr, D., Sanguinetti, G. & Grima, R. Validity conditions for moment closure approximations in stochastic chemical kinetics. The Journal of Chemical Physicsl 141, 084103 (2014).

106. Lakatos, E., Ale, A., Kirk, P. D. W. & Stumpf, M. P. H. Multivariate moment closure techniques for stochastic kinetic models. The Journal of Chemical Physics 143, 094107 (2015).

107. Lamperski, A., Ghusinga, K. R. & Singh, A. Stochastic optimal control using semidefinite programming for moment dynamics. Proc. of the 55th IEEE Conf. on Decision and Control, Las Vegas 1990–1995 (2016).

108. Ghusinga, K. R., Vargas-Garcia, C. A., Lamperski, A. & Singh, A. Exact lower and upper bounds on stationary moments in stochastic biochemical systems. Physical Biology 14, 04LT01 (2017).

109. Paulsson, J. Summing up the noise in gene networks. Nature (London) 427, 415–418 (2004).

110. Singh, A. & Bokes, P. Consequences of mRNA transport on stochastic variability in protein levels. Biophysical Journal 103, 1087–1096 (2012).

111. Gillespie, D. T. A general method for numerically simulating the stochastic time evolution of coupled chemical reactions. Journal of Computational Physics 22, 403–434 (1976).

112. Hooykaas, M. J. G. et al. Rna accessibility impacts potency of tough decoy microrna inhibitors. RNA Biology 15, 1410–1419 (2018).

113. Parra, M. et al. An important class of intron retention events in human erythroblasts is regulated by cryptic exons proposed to function as splicing decoys. RNA 24, 1255–1265 (2018).

114. Howard, J. M. et al. HNRNPA1 promotes recognition of splice site decoys by U2AF2 in vivo. Genome Research 28, 689–698 (2018).

115. Denichenko, P. et al. Specific inhibition of splicing factor activity by decoy RNA oligonucleotides. Nature Communications 10, 1590 (2019).

116. Padovan-Merhar, O. et al. Single mammalian cells compensate for differences in cellular volume and DNA copy number through independent global transcriptional mechanisms. Molecular Cell 58, 339–352 (2015).

117. Johnston, I. G. et al. Mitochondrial variability as a source of extrinsic cellular noise. PLOS Computational Biology 8, e1002416 (2012).

118. Shahrezaei, V., Ollivier, J. F. & Swain, P. S. Colored extrinsic fluctuations and stochastic gene expression. Molecular Systems Biology 4 (2008).

119. Hilfinger, A. & Paulsson, J. Separating intrinsic from extrinsic fluctuations in dynamic biological systems. Proceedings of the National Academy of Sciences 108, 12167–12172 (2011).

120. Singh, A. Transient changes in intercellular protein variability identify sources of noise in gene expression. Biophysical Journal 107, 2214–2220 (2014).

121. Shaffer, S. M. et al. Rare cell variability and drug-induced reprogramming as a mode of cancer drug resistance. Nature 546, 431–435 (2017).

122. Razooky, B. S., Pai, A., Aull, K., Rouzine, I. M. & Weinberger, L. S. A hardwired HIV latency program. Cell 160, 990–1001 (2015).

123. Chavez, L., Calvanese, V. & Verdin, E. HIV latency is established directly and early in both resting and activated primary CD4 T cells. PLOS Pathogens 11, e1004955 (2015).

124. Singh, A. & Weinberger, L. S. Stochastic gene expression as a molecular switch for viral latency. Current Opinion in Microbiology 12, 460–466 (2009).

125. Singh, A. Stochastic analysis of genetic feedback circuit controlling HIV cell-fate decision. Proc. of the 51st IEEE Conf. on Decision and Control, Maui, Hawaii 4918–4923 (2012).

126. Richman, D. D. et al. The challenge of finding a cure for HIV infection. Science 323, 1304–1307 (2009).

127. Dar, R. D., Hosmane, N. N., Arkin, M. R., Siliciano, R. F. & Weinberger, L. S. Screening for noise in gene expression identifies drug synergies. Science 344, 1392–1396 (2014).

